# Mutation of Ebola Virus Matrix Protein Cysteine Residues Increases Binding to Phosphatidylserine through Increased Flexibility of a Lipid Binding Loop

**DOI:** 10.1101/286120

**Authors:** Kristen A. Johnson, Nisha Bhattarai, Melissa R. Budicini, Carolyn M. Shirey, Sarah Catherine B. Baker, Bernard S. Gerstman, Prem P. Chapagain, Robert V. Stahelin

**Author notes:** Address Correspondence to: Robert V. Stahelin, Purdue University, 575 Stadium Mall Drive, West Lafayette, IN 47907.

## Abstract

The Ebola virus (EBOV) is a genetically simple negative sense RNA virus with only 7 genes yet it causes severe hemorrhagic fever in humans. The matrix protein VP40 of EBOV is the main driver of viral budding through binding to host plasma membrane lipids and formation of the filamentous, pleomorphic virus particles. To better understand this dynamic and complex process we have asked what the role of two highly conserved cysteine residues are in the C-terminal domain of VP40. Here we report that the mutation of Cys^311^ to alanine increases VP40 membrane binding affinity for phosphatidylserine containing membranes. C311A has a significant increase in binding to PS compared to WT, has longer virus like particles, and displays evidence of increased budding. C314A also has an increase in PS binding compared to WT, however to a lesser extent. The double Cys mutant shares the phenotypes of the single mutants with increased binding to PS. Computational studies demonstrate these Cys residues, Cys^311^ in particular, restrain a loop segment containing Lys residues that interact with the plasma membrane. Mutation of Cys^311^ promotes membrane binding loop flexibility, alters internal VP40 H-bonding, and increases PS binding. To the best of our knowledge, this is the first evidence of mutations that increase VP40 affinity for biological membranes and the length of EBOV virus like particles. Together, our findings indicate these residues are important for membrane dynamics at the plasma membrane via the interaction with phosphatidylserine.

## Introduction

The Ebola virus (EBOV) is a rare and neglected disease that causes hemorrhagic fever in humans with a high fatality rate. In 2014, the largest EBOV outbreak in history began. The outbreak became a global threat and over 3.6 billion dollars were spent to aid in containing the outbreak and treating patients (1). This outbreak highlighted the importance of gaining a basic understanding of how the EBOV components function, as there are still no FDA approved therapies for treatment or prevention of infection. While vaccine and antibody therapy against the EBOV glycoprotein has shown great promise, escape mutations against antibodies have been detected as EBOV was passaged through animals (2) and a watch list of potential glycoprotein mutations that may evade antibody control has been compiled (3).

With only 8 proteins expressed from its genome, EBOV is simple in design but is able to perform many functions through the transformative properties of the matrix protein, VP40 (4). VP40 is essential for viral success as it is key for transcriptional regulation during infection (4), formation of the lipid-enveloped virus particles, and virus particle budding (5,6).

Viral matrix protein flexibility is critical for multifunctional properties of VP40 and other viral matrix proteins (7). Through flexible regions and different conformations, VP40 is able to bind RNA, lipids, and interact with an abundance of host proteins to achieve transport to the plasma membrane and finally mediate viral budding (8–18). VP40 is sufficient form virus like particles (VLPs) that bud from cells independent of the other EBOV components (19).

VP40 binds to anionic lipids at the plasma membrane (PM) to form the host-derived lipid coat of the virus particle (20,21). VP40-lipid binding is also essential for the formation of VP40 hexamers (8,17,20) and the matrix sheet of large oligomers (8,17) VP40 interacts with the plasma membrane through its C-terminal domain (4,18,20,22) and forms oligomers through NTD-NTD interactions (18). When VP40 binds to anionic lipid phosphatidylserine (PS) hexamers are formed *in vitro* and in live cells (17,23). Lipid induced VP40 oligomerization allows dissociation of the N and C terminal domains that normally stabilize the protein in the closed dimer conformation. Experimental data and molecular dynamic simulations have shown that N- and C-terminal domain interactions, including several salt-bridges, are important for maintaining the closed conformation of the protein (18,23,24).

VP40 plasma membrane binding and localization has been found to be mediated by phosphatidylserine (PS) and the low abundance inositol lipid, PI(4,5)P_2_. Both lipids are critical to formation of VLPs where PS is important for initial plasma membrane recruitment and hexamer formation (17) and PI(4,5)P_2_ binding is essential for extensive VP40 oligomerization and oligomer stability (25). When lipid binding is abolished in cells, VP40 dependent budding is significantly reduced (17,20,22)

Cysteine is a rare and conserved amino acid with only 2.26% occurrence in mammals and higher eukaryotes, mathematically 0.865% less enriched than two codons should randomly produce (26). Cysteine residues are important for structural properties of proteins through disulfide bond formation and as a RNA, DNA, or histone binding protein through a zinc finger motif (27). More than 21% of cysteine residues are found in a C-(X)_2_-C motif, commonly attributed to metal binding and oxidoreductase proteins (26). Additionally, cysteine residues cannot engage in disulfide bond formation unless they have at least two residues between them (26).

VP40 has two highly conserved cysteine residues in the C-terminal domain of the protein. In many of the EBOV strains, these residues are found in a C-(X)_2_-C motif. In the crystal structure of VP40 from the Zaire type of EBOV (4LDB), the cysteine residues are not engaged in a disulfide bond (4). This is not surprising as VP40 is cytosolic and most cytosolic proteins do not have disulfide bonds as the reduced form is maintained by thioredoxin proteins (26). Because cysteine residues are low abundance residues, and VP40 has two cysteine residues in a C-(X)_2_-C motif, we hypothesized that they play an important role in VP40 function.

To test how these highly conserved residues may impact VP40 function we examined the lipid-binding and oligomerization properties of VP40 single or double cysteine to alanine mutations *in vitro* and in cells. Additionally, we measured VLP formation, filament length, and performed molecular dynamics simulations to investigate the molecular role of the Cys residues. Here we show the cysteine residues play an important role in regulation of PS binding and VP40 intramolecular interactions, loss of which increases VP40 affinity for PS and increases the length of filamentous virus like particles.

## Results

### VP40 has two adjacent cysteine residues in the C- terminal domain

VP40 cysteine residues are conserved in the Zaire, Tai Forrest, Bundibugyo, and Reston types but not the Sudan type of EBOV (**Figure 1A**). The Sudan VP40 also has two cysteine residues but they are located at slightly different positions, residues 314 and 320, respectively. **Figure 1B** shows the crystal structure (4LDB) with the two cysteine residues 311 (purple) and 314 (blue) highlighted. These residues are located within a solvent exposed, flexible loop on the side of the C-terminal domain of the protein. The cysteine residues are not resolved in the Sudan VP40 structure (4LD8) likely because they are also located within a flexible loop region of the protein. To investigate the role of these cysteine residues in VP40 function, mutation of each Cys to Ala was made in 6xHis-VP40-pET46 and EGFP-VP40 for bacterial expression and protein purification and expression in cell culture for imaging and budding experiments, respectively.

**Figure 1.**
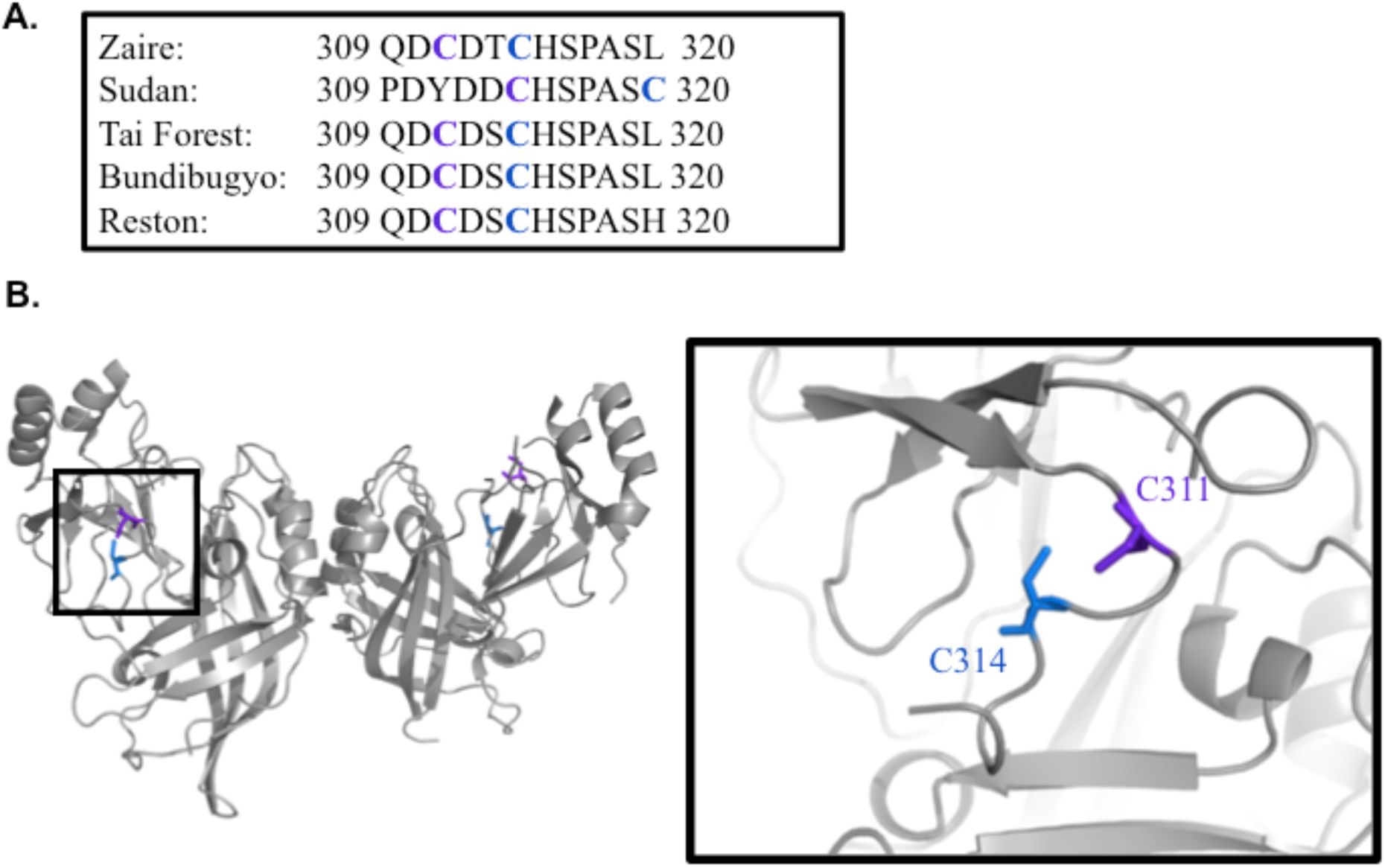
Ebola VP40 cysteine residues are conserved. **A**. With the exception of Sudan Ebola virus, cysteine at 311 and 314 are conserved. **B**. Structure of Zaire (4LDB) with C311 and C314 highlighted in the loop region.

### VP40 Cysteine Mutants have increased binding to Phosphatidylserine

VP40 binds to PS and PI(4,5)P_2_ in the plasma membrane of cells (17,25). To test if the cysteine residues have a role in lipid binding of VP40, VP40-WT, VP40-C311A, and VP40-C314A proteins were purified for lipid binding studies. Normally, the 6xHis-VP40-pET46 forms a dimer and octamer in solution and must be separated with size exclusion chromatography (See **Figure 5C**). The VP40 dimer was used in the binding studies here.

To test the lipid binding abilities of these mutants, a liposome pelleting assay was used. This assays allows for quantitative analysis of the fraction bound to control liposomes, PI(4,5)P_2_ containing liposomes, and phosphatidylserine (PS) containing liposomes (**Figure 3**). A representative SDS-PAGE is shown for each construct in **Figure 2A** where protein in the supernatant (SN) represents unbound protein and protein in the pellet (P) represents protein bound to the liposomes. The average fraction bound is shown in **Figure 2B**. C311A had a significant increase in binding to PS, while C314A a slight increase in PS binding compared to WT. C311A/C314A showed the most dramatic increase in binding to PS vesicles, an increase that was approximately the added effect of both single mutants on PS binding. Interestingly, there was no significant change in binding to PI(4,5)P_2_ in any of the cysteine mutants compared to WT (**Figure 2**). To the best of our knowledge, this is the first identification of VP40 mutations that increase the lipid-binding properties of this viral matrix protein.

**Figure 2.**
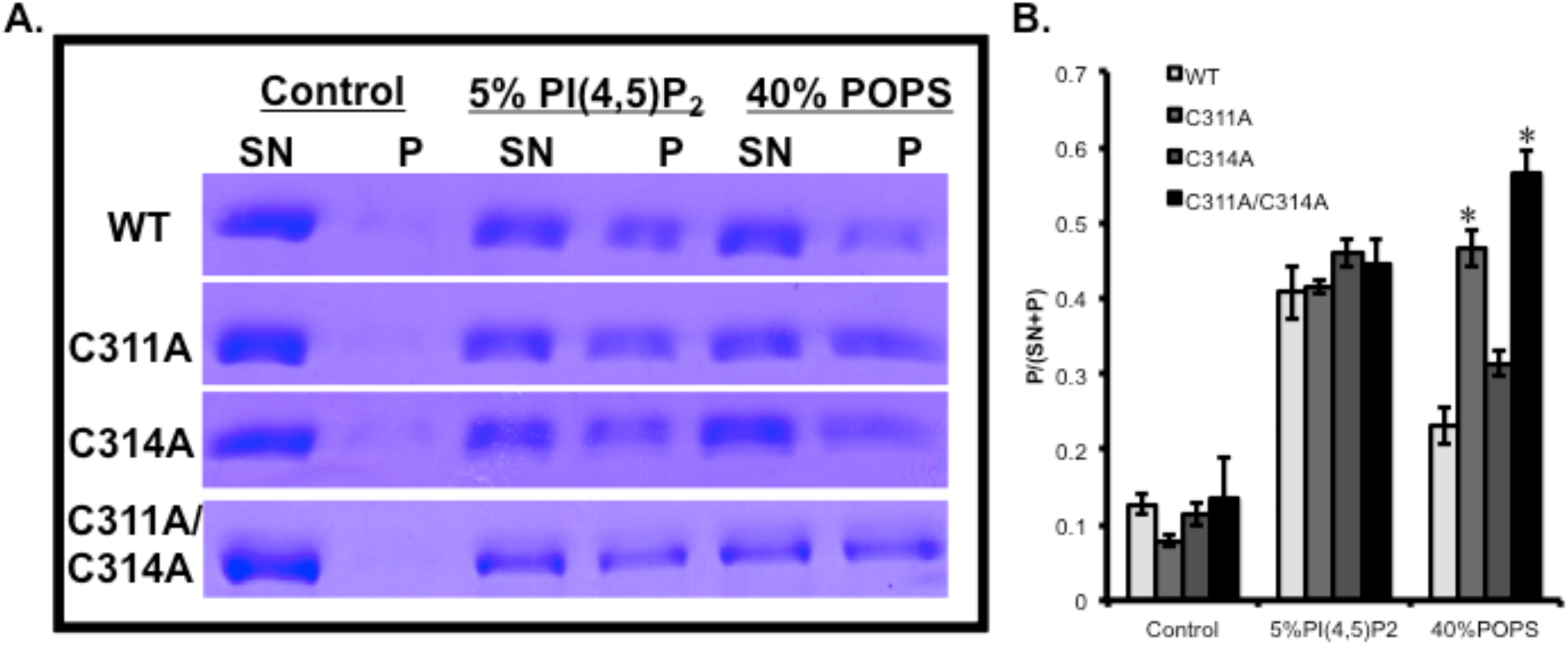
VP40 cysteine mutants have increased binding to the anionic lipid phosphatidylserine **A**. Representative SDS-PAGE of supernatant (SN) and pellet (P) samples from an LUV pelleting assay with VP40-WT, VP40-C311A, and VP40-C314A. **B**. Fraction of VP40 bound to lipids (P/(SN+P)) is shown as the average fraction bound over three replicates. Error bars represent the standard error of the mean, and * marks significance of a p value of 0.05 or less.

**Figure 3.**
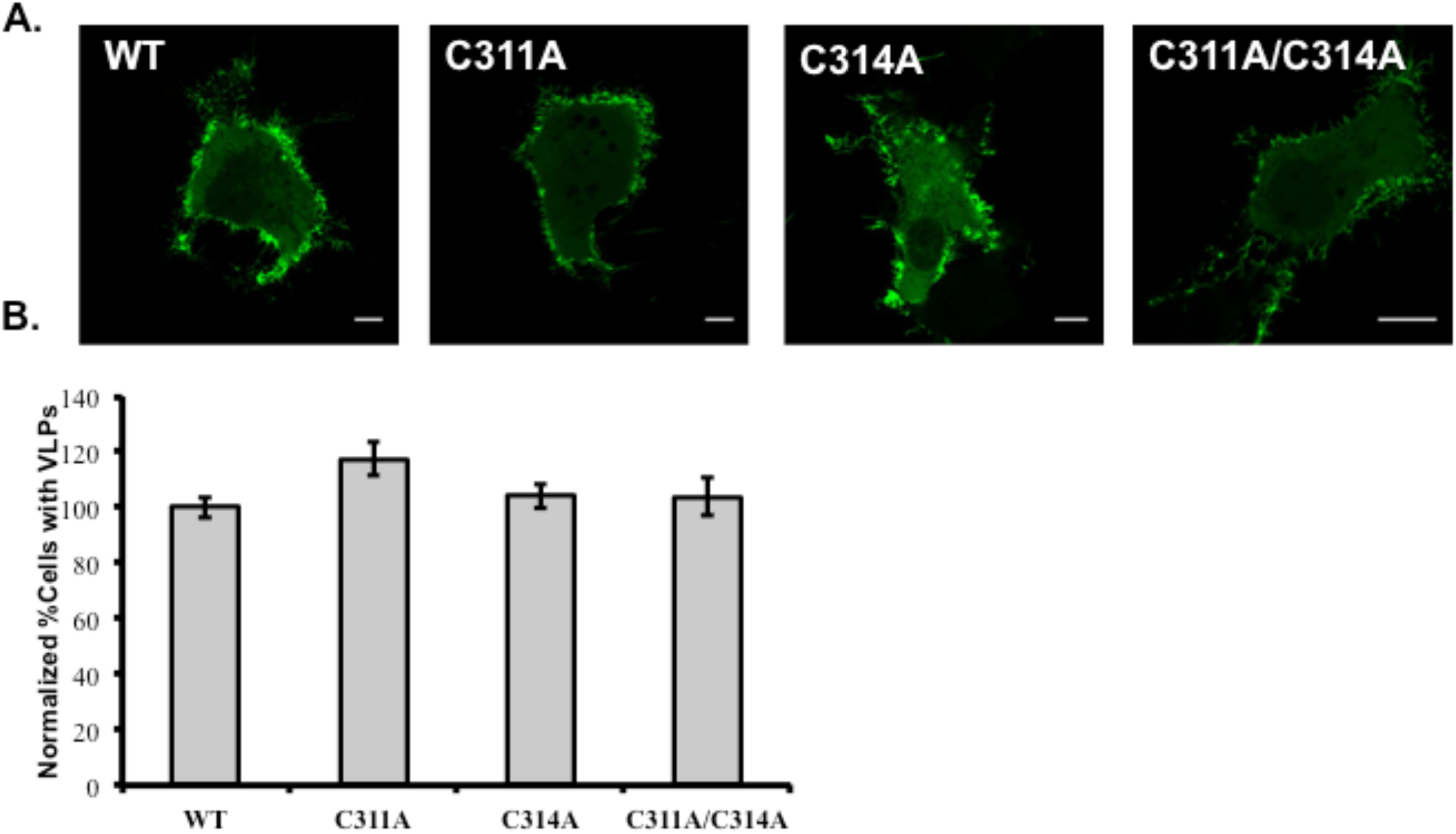
VP40 Cysteine mutants have normal phenotype when expressed in live cells. **A**. Representative images of EGFP-VP40-WT, EGFP-VP40-C311A, EGFP-VP40-C314A, and EGFP-VP40-C311A/C314A 12-14 hours post transfection in COS-7 cells. Scale bars are 10 µm. **B**. Cell populations expressing VLPs, data was normalized to EGFP-VP40-WT. Bars represent the average value over three experiments and error bars represent ± standard error of the mean.

### VP40 cysteine residues have minor effects on protein dynamics at the plasma membrane

Because VP40 cysteine mutants have an increase in binding to PS, an anionic lipid enriched in the inner leaflet of the plasma membrane of host cells that is critical for VP40 budding (17), we expressed the EGFP-tagged constructs in live cells to observe the phenotype of the mutants. EGFP-VP40-WT, EGFP-VP40-C311A, EGFP-VP40-C314A, and EGFP-VP40-C311A/C314A were expressed in COS-7 cells. Cells expressing VLPs or not were counted to see if the mutants had an altered ability to form virus like particles. Representative images of VP40-WT and mutants are shown in **Figure 3A**. While there was a slight increase in the number of cells containing VLPs emanating from the plasma membrane for C311A, this increase was not found to be statistically significant. Additionally, C314A and the double mutation exhibited similar VLP formation to WT VP40 where WT VLP formation was normalized to 100% (**Figure 3B**). The true population of cells with VLPs 14 hours post transfection is 70% with a standard deviation of 7.9% over three independent experiments.

Next, we investigated if there was a difference in VP40 membrane dynamics using fluorescence recovery after photobleaching (FRAP). The average normalized FRAP profile for each construct is shown in **Figure 4A** and the mobile fraction is shown in **Figure 4B**. C311A and C314A have a decreased recovery after photobleaching and a decreased mobile fraction, however, the decrease is not statistically significant in the single mutants (p=0.054 for C311A and p=0.175 for C314A). Double mutant C311A/C314A had a decreased mobile fraction from 0.36 in WT to 0.28 (p= 0.029). The diffusion coefficient was determined for WT (0.110 um^2^/s), C311A (0.108 um^2^/s) and C314A (0.135 um^2^/s) and C311A/C314A (0.104 um^2^/s) (**Figure 5C**). The results indicate the diffusion of WT and mutant VP40 constructs at the plasma membrane is within a similar range.

**Figure 4.**
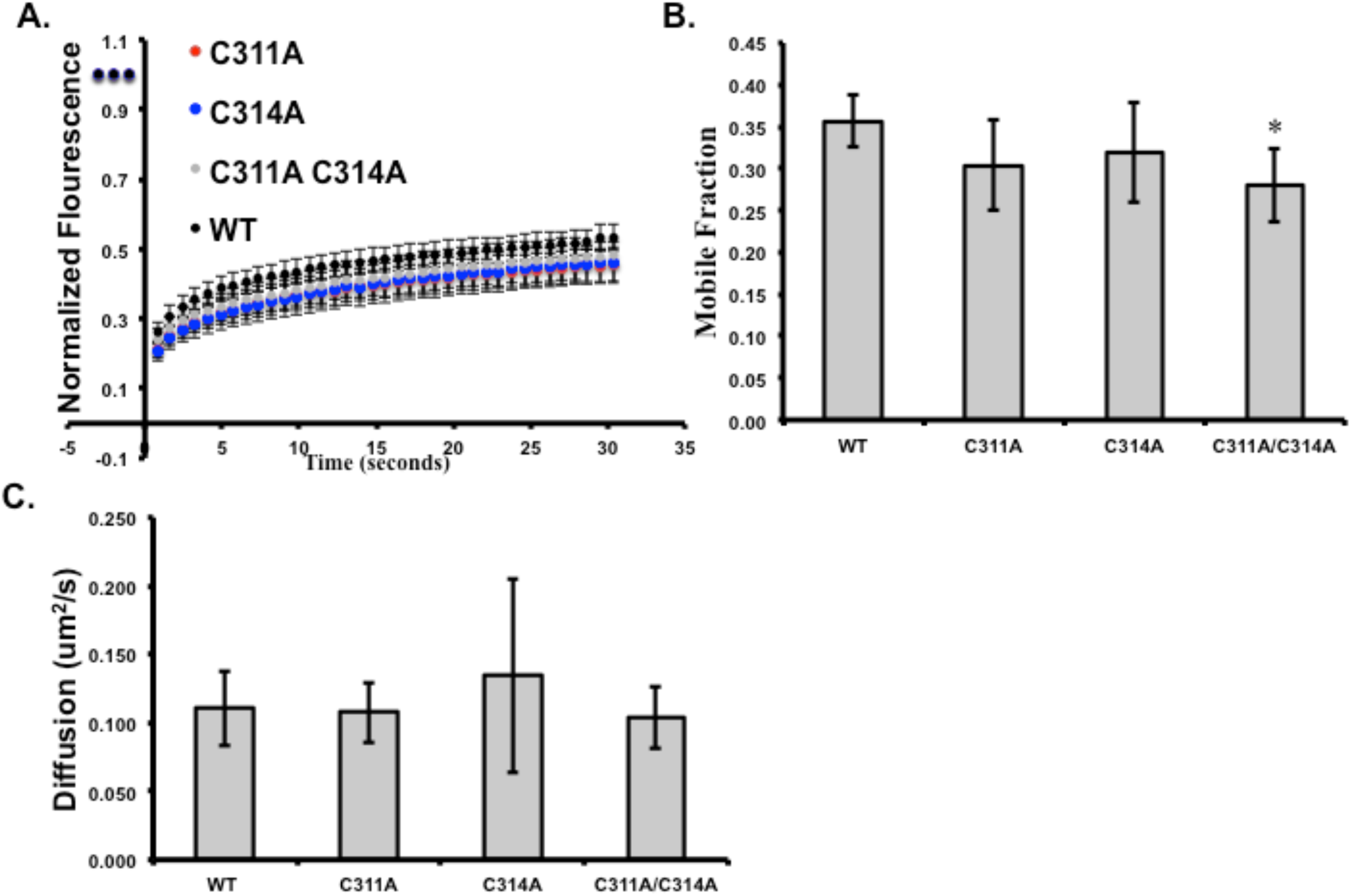
VP40 dynamics at the plasma membrane. **A**. Fluorescence recovery after photobleaching (FRAP) plot of EGFP-VP40-WT and mutants in COS-7 cells. **B**. The mobile fraction of EGFP-VP40-WT and mutants in COS-7 cells. **C**. The diffusion coefficient of EGFP-VP40-WT and mutants was determined and plotted. Values represent the average value over at least three experiments and error bars represent ± standard error of the mean. Significance is marked with a *, determined as a p value of 0.05 or less.

**Figure 5.**
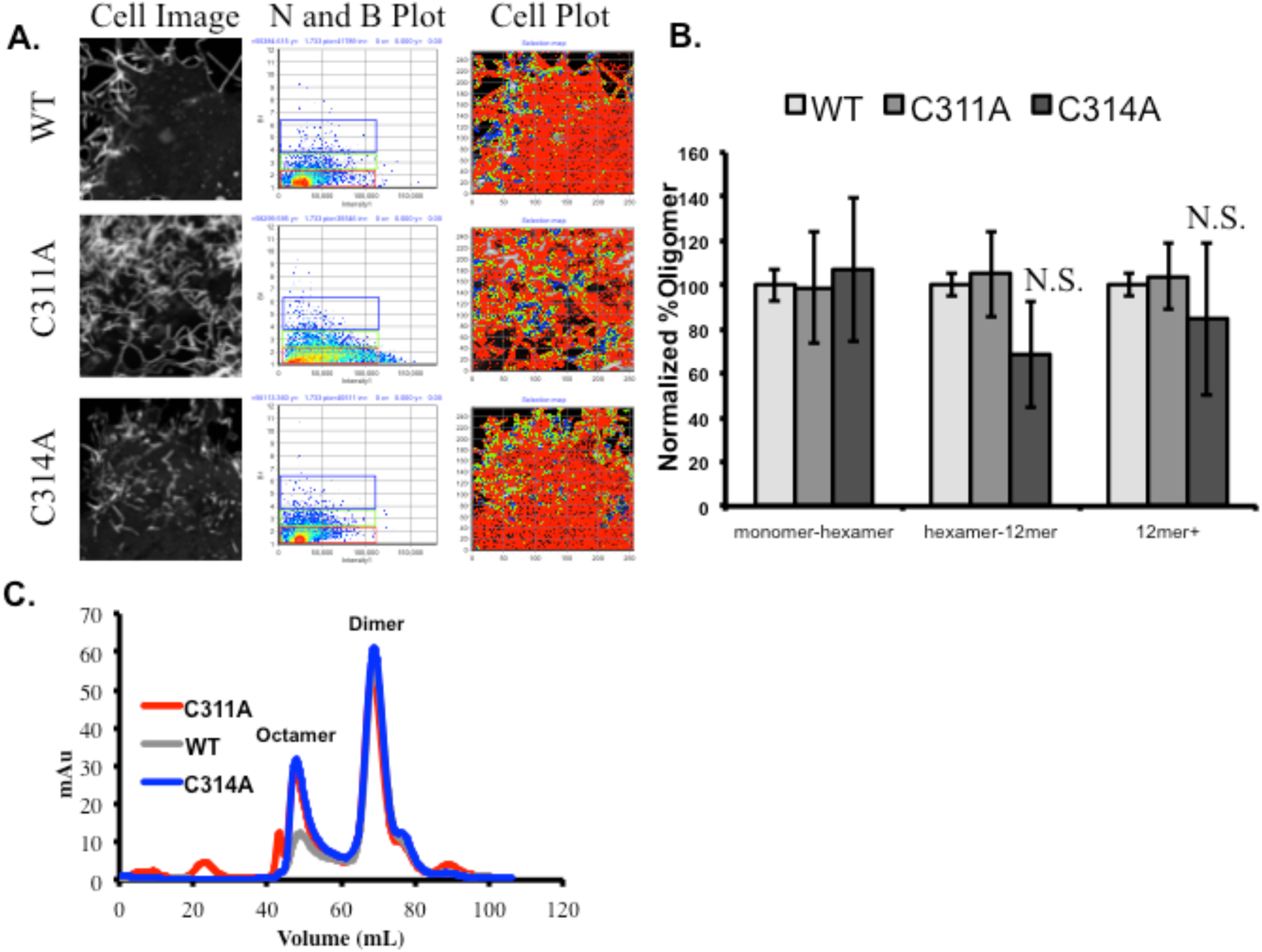
EGFP-VP40-WT and single mutant oligomer formation in cells and solution. **A**. Representative images from the number and brightness experiment to determine oligomer formation in live cells. **B**. Oligomer formation in cells normalized to EGFP-VP40-WT. No significant changes were observed. **C**. Size exclusion chromatogram trace of WT, C311A, and C314A purified VP40 protein shows similar formation of the octamer and dimer in solution compared to WT.

### Molecular dynamic simulations

#### Molecular dynamic studies demonstrate Cys residues regulate the position of a lipid-binding loop in VP40

We found that the Cys residues, especially Cys^311^, have an important role in C-terminal domain flexibility through interactions with a 199-G-SN-G-201 motif, which resembles a GxxG motif that has been found to be functionally important in other systems (28–30). Cys^311^ plays a role in a N-C terminal domain salt bridge as it is between the two CTD Aspartate residues (Asp^310^ and Asp^312^) that make strong ionic interactions with NTD residues R148 and R151 as shown in Fig. 6. Interdomain interactions between VP40 N- and C-terminal domain residues have been shown to be important in domain association, plasma membrane localization and VP40 oligomerization (24). Mutation of Cys to Ala altered the distance between the C_α_ atoms of residues 311 and residues 201. As shown in Figure 7B, the GSNG loop region extends toward the membrane during the simulation time, which helps to explain the enhanced PS binding by the C-terminal domain cationic residues in the C311A mutant as Lys residues near the GSNG motif have been proposed to play an important role in interacting with PS at the plasma membrane (4) and have been shown to be critical for interactions with PS containing vesicles (Del Vecchio et al. J. Biol. Chem., under review).

**Figure 6.**
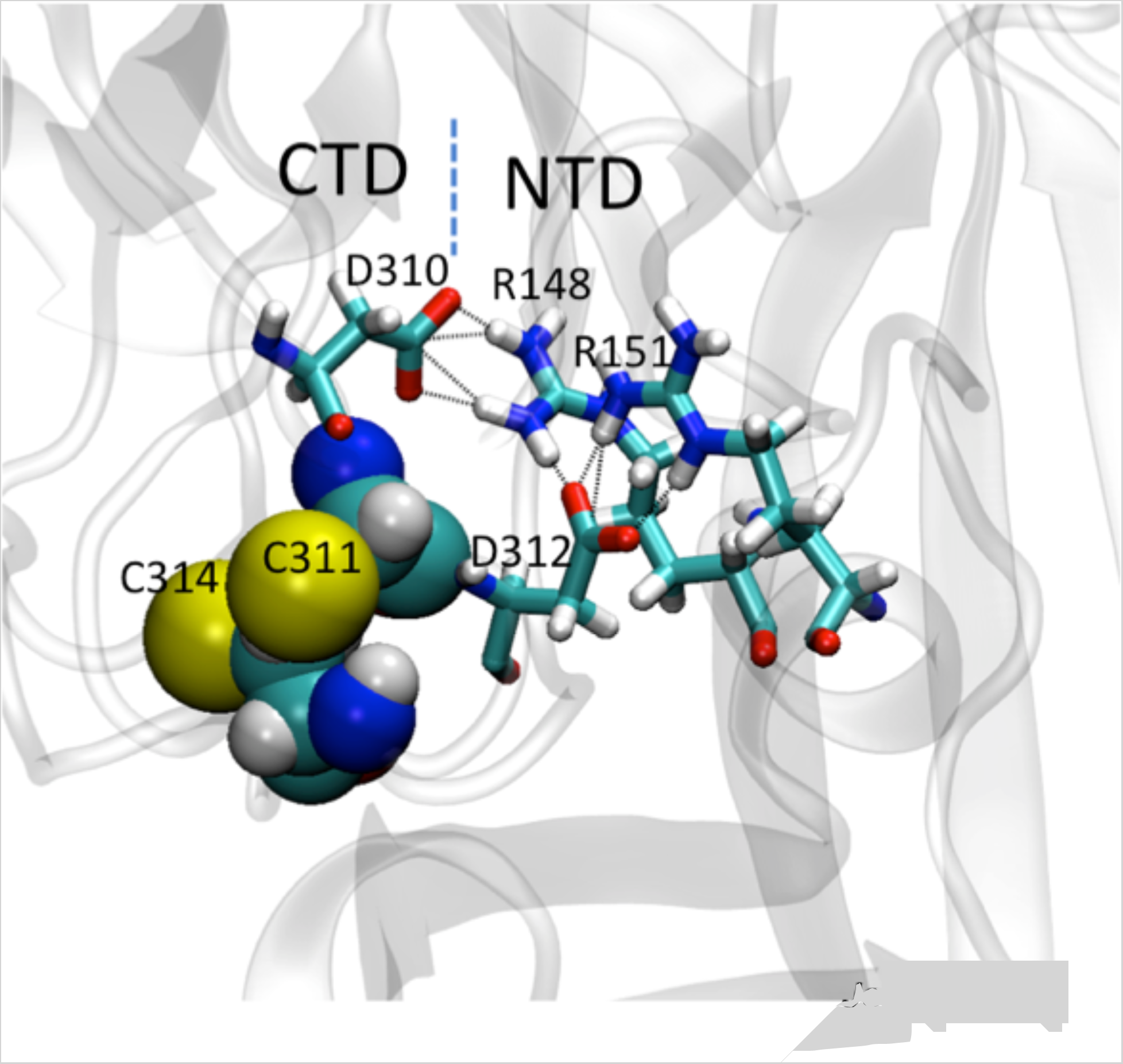
Aspartate residues 310 and 312 on either side of Cys^311^ are involved in the interdomain salt-bridges between N-and C-terminal domains.

**Figure 7.**
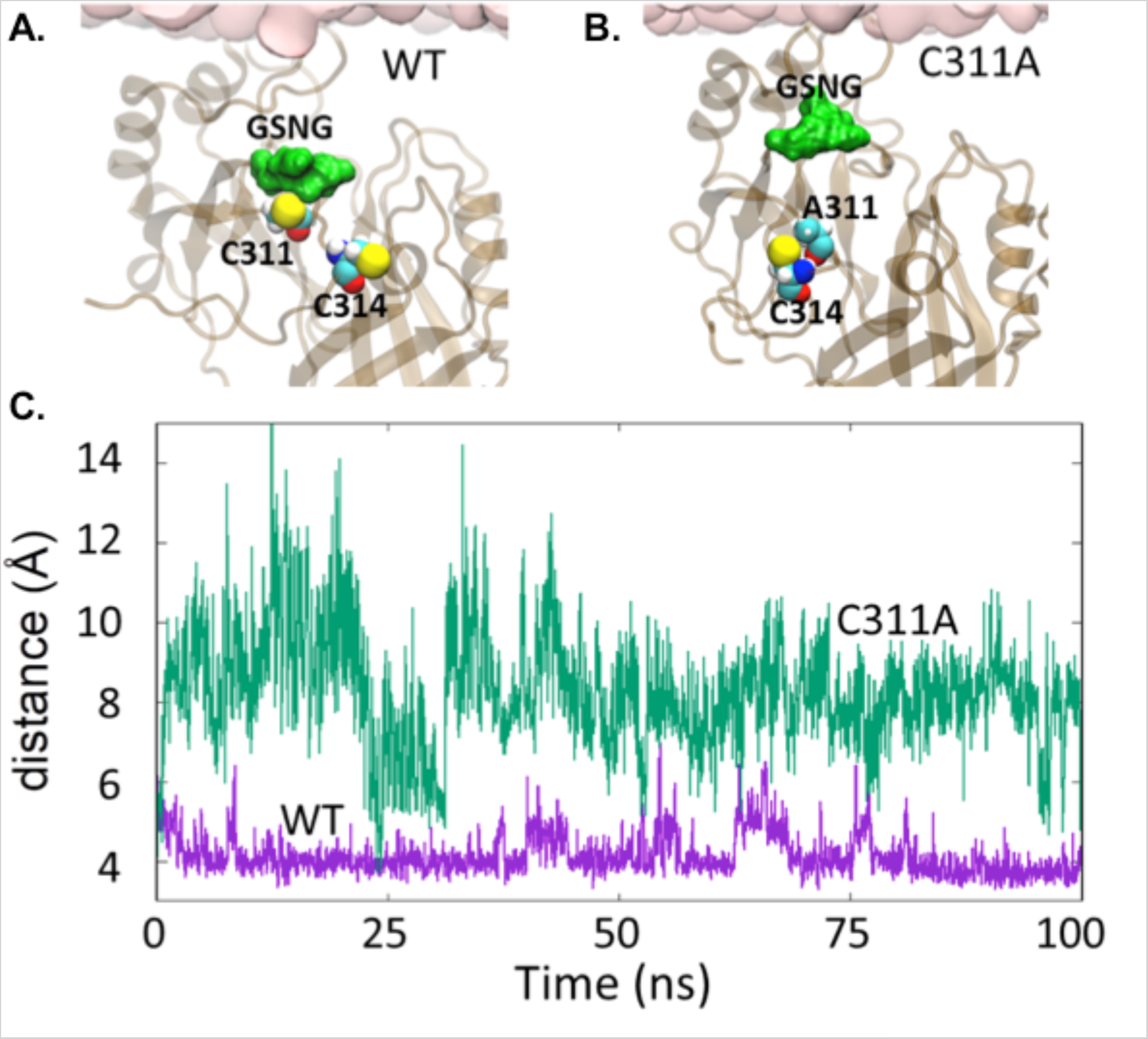
A. Cys^311^ interacting with the two polar residues (Ser^199^, Asn^200^) in the GSNG motif. B. Mutation C311A enhances the flexibility of the GSNG motif. Distance between the C_α_ atoms of residues 201 and 311 (purple: WT, green: C311A).

#### Mutation of VP40 Cys^311^ residue increases VLP filament length

When VP40 is expressed in mammalian cells, virus like particles form at the surface of the plasma membrane. The particles can be observed using various imaging techniques including confocal microcopy with fluorescently tagged VP40 or scanning electron microscopy (SEM) to visualize structures on the surface of a cell. We have utilized SEM in the past to visualize VLP production by VP40 in COS-7 cells and found a similar phenotype to EBOV expressed in this cell line (31). We used SEM to visualize the VLPs on the surface of the WT and single mutant transfected cells (**Figure 8A**). The VLP length was measured in FIJI and the values were plotted as a histogram to show the distribution of particle length. C311A had a larger population of 3 µm and longer VLPs than WT and C311A (**Figure 8B**).

**Figure 8.**
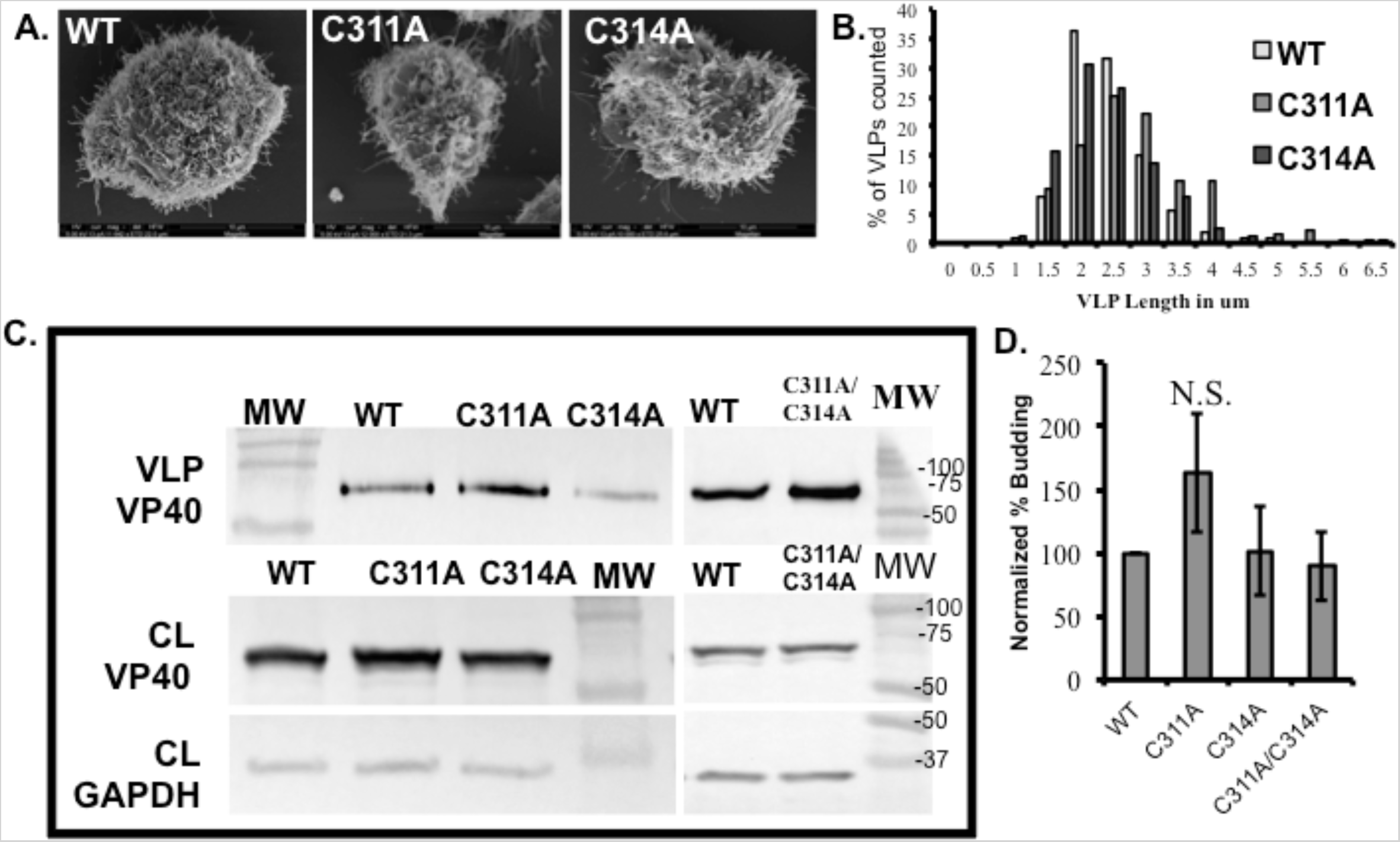
VP40 VLP production and budding efficiency is not significantly changed without cysteine residues. **A**. Representative scanning electron micrographs of EGFP-VP40-WT, and single mutants to show VLPs on the surface of the cell. **B**. Histogram of VLP length for EGFP-VP40-WT and single mutants, values are represented as the percentage of the population of VLPs at each length bin for each construct. **C**. Western blot of VP40 in the cell lysate (CL) and VLP samples. GAPDH was used as the loading control for the CL samples. **D**. Normalized budding efficiency for EGFP-VP40-WT and single Cys mutants. Average values are normalized to WT budding efficiency and represent at least three experiments. Error bars are ± standard error of the mean.

To determine if the increase in PS binding by C311A or C314A would affect VP40 budding efficiency, VLPs and cell lysates were collected 24 hours post transfection. Normally, 24 hours post transfection, an abundance of VLPs have been released from the cell and can be easily detected with immunoblotting or ELISA. Through analysis of VP40 in the VLP and CL samples, budding efficiency can be determined. In this study we used western blotting to determine the budding efficiency of Cys mutants compared to WT VP40 (see methods for details). GAPDH, an enzyme involved in glycolysis, was used as the loading control. VP40-C311A (N=4) had an increase in VLP budding efficiency compared to WT (N=7) while C314A (N=4) and C311A/C314A (N=3) had a similar budding efficiency (**Figure 8C and 8D**). While the increase in VLP formation for C311A compared to WT was detectable, it should be noted the increase wasn’t statistically significant. Nonetheless, the increased PS binding affinity and filament length, combined with computational evidence of these Cys residues regulating the position of the membrane binding loop, underscores the importance of examining individual mutations of VP40 in assembly, budding and transcriptional regulation of EBOV.

## Discussion

VP40 lipid binding properties are essential for efficient budding from the plasma membrane of mammalian cells (17,25). In this study, we found that the VP40-cysteine residues play an important role in how VP40 interacts with PS containing membranes. When VP40 Cys^311^ was mutated to alanine in the single or double mutant, there was a significant increase in binding to PS. The increase in binding to PS in C314A was of lesser magnitude and not statistically significant compared to WT PS binding. Molecular dynamic simulations provided mechanistic details to this observation. For instance, Cys^311^ plays a predominant role in interacting with the 198-G-SN-G-201 motif, which is adjacent to the membrane interacting loop residues Lys^224^ and Lys^225^, which has been suggested to be key in binding the plasma membrane (4) and are critical residues to binding PS *in vitro* (Del Vecchio and Stahelin, *J. Biol. Chem*. under review). The polar residues Ser and Asn in the motif interact with Cys^311^ playing a somewhat restrictive role in this loop regions ability to extend towards the membrane surface. As shown in Fig. 7, the GSNG motif interacts with Cys^311^ (Fig. 7A), whereas it is relatively unrestrained and distant from A311 (Fig. 7B) in the C311A mutant. This is reflected in the C_α_− C_α_ distance between Cys or Ala (residue 311) and Gly^201^ of GSNG loop (Fig. 7C), which shows a relatively steady loop position for the WT. Interestingly, this loop is found to open and close intermittently in the absence of membrane interactions as shown in the movie S1.

Previous research by Adu-Gyamfi et al. identified that PS becomes exposed on the exterior of VP40 VLPs following accumulation of VP40 at the plasma membrane inner leaflet (17). The PS content of the plasma membrane is also a critical component of EBOV assembly and budding as a 35-40% reduction in plasma membrane PS was sufficient to significantly reduce VLP formation and detectable sites of VP40 budding (17). However, it is unknown if increasing the plasma membrane PS content or VP40 affinity for PS would increase the amount of VLPs formed. The budding increase in the C311A mutant was dramatic but not significant and changes in C314A and the double mutant were very similar to WT suggesting that C314A may reduce the effect of C311A in cells in the double mutant construct.

The increased filament length of VLPs for C311A was consistently and significantly above WT VLP length. While still an unknown area of research, increased VLP length could be attributed to an increase in PS binding or altered inter- or intradomain contacts by VP40. Throughout the 2014-2016 EBOV outbreak as well as passage of the EBOV through animal studies, several mutations of VP40 have been found (32,33). For the most part, the functional consequences of these mutations are unknown but it is feasible to hypothesize mutation of Cys^311^, its interacting residues or residues adjacent to this region, could increase VP40 affinity for the plasma membrane PS in the authentic virus.

Some cysteine residues or motifs containing multiple Cys residues are important for nucleotide binding (27). VP40 is known to form an octameric ring and can bind RNA with N-terminal domain residues (4,34). There was no difference detected in VP40 oligomers in cells (Figure 5A and B) and the cysteine mutants had a normal fraction of octamer compared to the WT protein when purified from *E. coli* (**Figure 5C**). Number and Brightness analysis was used to determine oligomerization of each VP40 construct at the surface of the cell. There were no significant differences in VP40 oligomerization with WT, C311A, or C314A after three independent experiments. Representative images with corresponding N&B plots and cell plots are shown in **Figure 5A**. The population of monomer-hexamer, hexamer-12mer, and 12mer+ were normalized to the value of WT and plotted ± the standard error of the mean (**Figure 2B**). When VP40 is purified from bacteria, octamer and dimer populations are observed. Because these two populations have different structures and functions in the context of the virus, they must be separated for binding studies to better understand VP40 lipid binding. VP40-C311A and C314A have similar profiles to WT VP40 indicating there are no major structural changes with these mutants. A similar profile was also observed for the double mutant (data not shown). Additionally, there was not an obvious change in localization or accumulation of EGFP-VP40-WT or Cys mutants in the nucleus or perinuclear region (confocal imaging, **Figure 3**). While these results are consistent across *in vitro* and cellular experiments, future research is required to understand if these Cys residues have a role in transcriptional regulation in the context of the live virus.

In HIV studies, one group found that when cys residues in P7 were mutated, it was unable to bind nucleotides. While it couldn’t bind nucleotides, a similar budding compared to WT virus was observed. However, the virons produced were non-infectious (35). The role of the VP40 transcriptional regulation is not completely understood. In 2013, Bornholdt and colleagues discovered that VP40 reduces transcription by approximately 70% (4). The RNA binding mutant, R134A, only reduced transcription approximately 30% (4). While nucleotide binding properties were lost in R134A, this mutant retained its budding properties (4). If Cys^311^ and/or Cys^314^ are important for nucleotide binding or transcriptional regulation via a zinc finger motif, the mutants could be considered consistent with what was found with the R134A phenotype and respective budding compared to WT VP40. Additionally, the CTD residues were not previously implicated in nucleotide binding in the structural studies of the octameric ring. In fact, the CTD is highly flexible and not resolved in octameric ring structures available. Nonetheless, further research is needed to investigate potential metal binding and/or nucleotide binding abilities of VP40’s C-(X)_2_-C motif.

### Experimental Procedures

#### Molecular Biology

Site directed mutagenesis was performed with the QuikChange II XL Site-Directed Mutagenesis Kit according to the manufacturer’s instructions (Agilent Technologies, Santa Clara, CA). Primers were ordered from Integrated DNA Technologies (Coralville, IA) according the specifications in the QuickChange kit. Primers used were (listed 5’ to 3’): C311A Forward:C ACA CAG GAT GCT GAC ACG TGT CAT TCT CCT GC, Reverse: GC AGG AGA ATG ACA CGT GTC AGC ATC CTG TGT G. C314A Forward: C ACA CAG GAT TGT GAC ACG GCT CAT TCT CCT GC, Reverse: GC AGG AGA ATG AGC CGT GTC ACA ATC CTG TGT G. C311A/C314A (single mutant as template DNA) Forward: C ACA CAG GAT GCT GAC ACG GCT CAT TCT CCT GC, Reverse: GC AGG AGA ATG AGC CGT GTC AGC ATC CTG TGT G. Mutations were sequenced at the Notre Dame sequencing facility with sanger sequencing. Sequencing files were analyzed with 4Peaks.

#### Protein Expression and Purification

6xHis-VP40 pET46 was transformed into Rosetta BL21 DE3 cells according to the manufacturer’s instructions. Transformed bacteria were grown at 37°C until an OD_600_ of 0.6-0.9 was achieved. Protein expression was induced with 1 mM IPTG at room temperature for 4-6 hours. Bacteria were pelleted and stored at -20°C before purification. Protein purification was performed as previously described in detail (25). Eluted protein was further purified using size exclusion chromatography to isolate the VP40 dimer from the dimer-octamer mixture. Protein concentration was determined with a BCA assay and stored at 4°C in 10 mM TRIS, pH 8.0 containing 300 mM NaCl for up to 2 weeks.

#### Liposome Pelleting Assay

The liposome compositions used were as follows: control liposomes (DPPC: Chol: dansylPE 49:49:2), PI(4,5)P_2_ containing liposomes (DPPC: Chol: PI(4,5)P_2_: dansylPE 46.5:46.5:5:2), and PS containing liposomes (DPPC: Chol: POPS: dansylPE 29:29:40:2). Lipids were dried under a nitrogen gas stream and stored at -20°C until use. Lipid films were hydrated with 250 mM raffinose pentahydrate in 10 mM Tris, pH 7.4 containing 150 mM NaCl. Liposomes were extruded through a 200 nm filter and dynamic light scattering was used to check size. Once size was verified, raffinose liposomes were diluted and added to the protein/buffer mixture for the binding experiment. Protein and liposomes were incubated for 30 minutes at room temperature then centrifuged to pellet liposomes (75,000 *x g* for 30 minutes at 22 °C). Supernatants (SN) were removed and pellets (P) resuspended under a UV wand to visualize the pellet (2% dansylPE is included in the liposomes). SN and P fractions were run on SDS PAGE and protein density was determined using ImageJ. All liposomes were used the same day they were made and the raffinose buffer was made fresh each day.

#### Molecular Dynamics Simulations

The crystal structure of the VP40 dimer was taken from the Protein Data Bank (PDB ID: 4LDB). The protein and plasma membrane (PM) complex was set up using *Charmm-Gui* web server (36,37). The PM for the wild-type contained 1-palmitoyl-2-oleoyl-sn-phosphatidylcholine (POPC), 1-palmitoyl-2-oleoyl-sn-phosphatidylethanolamine (POPE), 1-palmitoyl-2-oleoyl-sn-phosphatidyl-serine (POPS), palmitoylsphingomyelin (PSM), palmitoyl-oleoyl-phosphatidyl-inositol (POPI) and cholesterol (CHOL). The lipid composition in the lower leaflet of the PM was 11:32:17:9:10:21 (POPC:POPE:POPS:POPI:PSM:CHOL). There were 149 lipids on the upper leaflet and 151 on the lower leaflet. The system was solvated using TIP3 water molecules and neutralized with counter ions. The total system consists of ∼122,000 atoms. The system for mutant C311A was set up with similar lipid ratios.

All-atom molecular dynamics simulations were performed using NAMD 2.12 (38) and using CHARMM36 force field (39). The particle mesh Ewald (PME) method (40) was used to calculate long range electrostatic interaction and SHAKE algorithm was employed to constrain the covalent bonds. The system was minimized for 10,000 steps with six steps equilibration given by *Charmm-gui* (37). A Nose-Hoover Langevin-piston was used to control the pressure with piston period of 50 fs decay and 25 fs. The Langevin temperature coupling with friction coefficient of 1 ps^−1^ was used to control the temperature. All production runs were performed with a 2 fs time step. Visualization and rendering were done with Visual Molecular Dynamics (VMD) (41).

#### Cell culture and transfection

COS-7 cells were maintained and transfected as previously described in detail (25).

#### Confocal Imaging

A Zeiss 710 LSCM was used to visualize EGFP-VP40 phenotypes in live cells. Number and Brightness data acquisition was performed on an Olympus FV2000 microscope and data analysis was performed with SimFCS as described previously (42–44).

#### Fluorescence recovery after photobleaching (FRAP)

FRAP was performed as described previously with some modifications (45). A 2.2 μm ROI was used to bleach EGFP with 50 iterations of 100% 488 nm laser power after three pre-bleach scans. Fluorescence recovery was measured for 30 seconds following the beaching. Data represents 20 images per construct collected over two different days and 4 independent experiments.

#### Scanning Electron Microscopy

COS-7 cells were transfected for 12-16 hours then scraped from plates, pelleted through centrifugation (900 *rcf*, 6 min), washed with 1X PBS then suspended in primary fixative. SEM processing was performed as previously described in great detail (25).

#### VLP collection

VLP collections were performed 24 hours post transfection as previously described (25). Briefly, the supernatant of the transfected cells was collected and applied to a 20% sucrose cushion. Samples were centrifuged at 100,000 x *g* for 2 hours. VLP pellets were suspended in 150 mM ammonia bicarbonate. Cells were trypsinized from the plate and lysed with RIPA buffer (with protease inhibitors) for 1 hour on ice with intermediate vortexing. Samples were centrifuged at 25,000 x *g* for 17 minutes and the soluble fraction was removed. A BCA was used to determine the cell lysate (CL) protein concentration for analysis with western blotting.

##### Western Blot

CL and VLP samples were analyzed via western blotting to determine the budding efficiency of WT VP40 and respective mutations. 15 μg of protein was loaded for the CL samples while 5x volume of CL samples were run for VLP samples. For VP40 detection the following antibodies were used (F56-6A1.2.3, ThermoFisher Scientific) primary antibody and HRP (AB 6808, Abcam, Cambridge, United Kingdom) secondary. For GAPDH detection, (AB 8245, Abcam) primary and HRP (AB 6808, Abcam) secondary were used. ECL detection was used to visualize VP40 and GAPDH according to the manufacturer’s instructions (ThermoFisher). The percent budding was determined with analysis in FIJI (46). Blots are representative of at least three independent experiments.

## Acknowledgements

**These studies were supported by the NIH (AI081077) and the Indiana University School of Medicine-South Bend Imaging and Flow Cytometry core to R.V.S**. K.A.J would like to thank Tatyana Orlova for assistance in scanning electron microscopy. K.A.J and R.V.S would like to thank Jessica Brown for helpful discussions.

## Author Contributions

K.A.J, R.V.S and C.M.S planned and designed the experiments and K.A.J, M.R.B, C.M.S, and S.C.B performed experiments. K.A.J and C.M.S performed the data analysis. N.B., B.S.G, and P.P.C. designed and performed molecular dynamics simulations. K.A.J and R.V.S wrote the manuscript. All authors reviewed the results and approved the final version of the text.

